# Computationally-guided design and selection of ribosomal active site mutants with high activity

**DOI:** 10.1101/2022.06.02.493746

**Authors:** Camila Kofman, Andrew M. Watkins, Do Soon Kim, Alexandra C. Wooldredge, Ashty S. Karim, Rhiju Das, Michael C. Jewett

## Abstract

Understanding how modifications to the ribosome affect function has implications for studying ribosome biogenesis, building minimal cells, and repurposing ribosomes for synthetic biology. However, efforts to design sequence-modified ribosomes have been limited because point mutations in the ribosomal RNA (rRNA), especially in the catalytic active site (peptidyl transferase center; PTC), are often functionally detrimental. Moreover, methods for directed evolution of rRNA are constrained by practical considerations (e.g., library size). Here, to address these limitations, we developed a computational rRNA design approach for screening guided libraries of mutant ribosomes. Our method includes *in silico* library design and selection using a Rosetta stepwise Monte Carlo method (SWM), library construction and *in vitro* testing, and functional characterization *in vivo*. As a model, we apply our method to making modified ribosomes with mutant PTCs. We engineer ribosomes with as many as 30 mutations in their PTCs, highlighting previously unidentified epistatic interactions, and show that SWM helps identify sequences with beneficial phenotypes as compared to random library sequences. We further demonstrate that some variants improve cell growth *in vivo*, relative to wild type ribosomes. We anticipate that SWM design may serve as a powerful tool for high-resolution rRNA design.

## Introduction

The ribosome is a complex macromolecular machine that has evolved to synthesize proteins by catalyzing peptide bonds between amino acids. Essential for all life, the ribosome is considered to be a ribozyme, as its catalytic active site, the peptidyltransferase center (PTC), is primarily composed of ribosomal RNA (rRNA) (1–4). Consequently, ribosome assembly and function are tightly linked to RNA folding and stability, and the ribosome’s structure has been evolutionarily constrained to maintain a free energy minimum that confers the ability to form peptidyl bonds rapidly and reliably (5, 6).

The production of modified ribosomes with biochemical defects (7) or altered capabilities (e.g., β-amino acid incorporation (8, 9)) serves as an important tool to better understand molecular translation and enable synthetic biology applications (10). However, designing and engineering mutant ribosomes is far from a trivial undertaking because ribosomal mutations can disrupt translation in ways that are often lethal to cells. For example, previous efforts to create modified ribosomes have shown that making even single point mutations to the PTC sequence can nullify the ribosome’s ability to properly assemble or catalyze bond formation (11–14). Not only is it difficult to identify functional small-scale mutations, but some detrimental mutations can also be rescued by synergistic mutations in adjacent or distal regions, highlighting the design challenges of rRNA engineering and importance of considering epistatic relationships between residues when designing libraries (15, 16).

While tethered ribosomes (17–20) or cell-free strategies (21–25) can be used to identify functionally detrimental ribosomal mutations, efforts to build modified ribosomes remain hampered by practical considerations in making and evaluating rRNA libraries. For example, the combinatorial space for rRNA evolution is large, such that random mutagenesis and selection approaches cannot be feasibly used to screen all possible variants. In addition, due to primer bias, randomized libraries constructed using PCR are known to have imbalanced initial populations, skewing assessments of a library’s members by overemphasizing the more common ones (26). PCR-based library construction approaches are also difficult to apply to multiple regions of rRNA that are close in three-dimensional space but not primary sequence space, which is common in the structurally complex PTC (27). A further challenge is that DNA libraries are typically propagated in cells, where transformation idiosyncrasies limit library size (8, 28–30). Alternative approaches are thus needed to test unbiased rRNA libraries of larger size and complexity such that we can explore diverse energy landscapes via large-scale sequence changes and identify mutant ribosomes with significantly modified architectures.

Here, to explore rRNA design rules with high-throughput methods for identifying synergistic mutations, we develop a computationally guided approach for making modified ribosomes. First, we use a stepwise Monte Carlo method (SWM) in Rosetta to score, rank, and select rRNA library members using an all-atom energy score (31) in an unbiased way. While previous methods have used computationally expensive, low-resolution coarse graining or small perturbations to fully build conformations (32), SWM requires much less computational power to reach an equivalent level of atomic accuracy. This approach also allows us to define libraries in three-dimensional space, including any residues that are potentially interacting and could mutate to play compensatory roles. We combine this computational approach with a high throughput *in vitro* ribosome synthesis, assembly, and translation (iSAT) screening platform (10, 33–35) to test the computationally identified mutants, allowing us to rapidly assay promising candidates. The resulting ribosome mutants highlight the surprising flexibility of the PTC to large-scale mutations and elucidate previously unknown epistatic relationships between distal regions of the PTC.

Unexpectedly, many of these highly mutated variants can support life in cells with only minor phenotypic effects. We anticipate that our high-throughput, computationally guided approach will allow for improved studies of complex rRNA libraries to ultimately enable novel ribosomal activity as well as deeper understanding of molecular translation.

## Materials & Methods

### SWM design simulations

To obtain an initial structure for stepwise Monte Carlo design, a crystal structure of the *Escherichia coli* ribosome (PDB code: 4YBB) was obtained and loaded into PyMOL. The residues of interest, local to a particular site in the ribosome, were selected, and that selection was expanded to include a 25.0 Å sphere of neighboring residues, enough to encompass several shells of indirect interactions. The full selection, including both residues of interest and neighbors, were saved to a ‘native’ PDB file. This ‘native’ file was passed to a Python script distributed with the Rosetta application, *tools/rna_tools/pdb_util/pdb2fasta*.*py*, to obtain a corresponding FASTA-formatted file with appropriate numbering. Finally, the 25.0 Å sphere of neighbors, but omitting the actual residues of interest, were saved to a ‘starting’ PDB file, ready for stepwise Monte Carlo design.

The sequence positions within the FASTA file that corresponded to the residues of interest were edited with a text editor to ensure the design simulation would sample any nucleotide but the wild-type nucleic acid identity: *a* was changed to *b* (the IUPAC ambiguous single-letter code representing ‘anything but adenosine’); *c* was changed to *d*; *g* was changed to *h*; *u* was changed to *v*. Because the region being redesigned would be free to resample its backbone conformation, some ‘adaptation’ between the fully flexible designed region and the totally rigid crystal context was necessary. To this end, the residues adjacent in primary sequence to any redesigned residue were indicated to the *-extra_min_res* flag: these residues, although not subject to explicit backbone sampling, were subject to quasi-Newtonian energy minimization along with the designed residues during simulation.

Simulations for H75, H73, and H91 were run for 1000 Monte Carlo cycles, while simulations for H92 were run for 2000 Monte Carlo cycles due to its structural complexity. At least 10,000 independent trajectories were run for each library. Full code examples for setting up, conducting, and analyzing design simulations are provided at https://github.com/everyday847/ptc_swm_modeling; documentation for stepwise Monte Carlo that includes details on design simulations is available at https://new.rosettacommons.org/docs/latest/application_documentation/stepwise/stepwise_monte_carlo/stepwise.

### Forward folding with SWM

In a design simulation, different sequences may be the lowest scoring frame of a trajectory at significantly variable frequencies. As a result, it can be difficult to make confident comparisons between the best energy sampled for two sequences or to set a strict threshold to select a small number of desired variants. Instead of establishing a strict cutoff selecting only a few variants for experimental characterization based on variable quantities of data, we ran individual ‘forward-folding’ simulations on a larger number of fixed sequences, using the 200 top-scoring sequences from the design simulation. These ‘forward-folding’ simulations used only 500 cycles and generated exactly 400 models each, ensuring an ‘apples to apples’ comparison among sequences for the final selection that would be inaccessible to a design simulation alone. We ran these simulations specifically for H75, where we were interested in whether lower scores would correlate to superior activity so accuracy and fair sampling for the single highest score was paramount. We elected not to repeat the simulations for the other helices due to computational resource constraints. Full code examples for setting up, conducting, and analyzing forward folding simulations are provided at https://github.com/everyday847/ptc_swm_modeling.

### Plasmid construction & preparation

Plasmids were ordered from Twist in two backbones: one in the pT7rrnB backbone (34) and one in pAM552G (19). Plasmids used for testing of variants in iSAT were built using pT7rrnB, a 7,311-bp plasmid. This plasmid carries an *E. coli* rRNA operon, rrnB, under the control of the T7 promoter, as well as the ampicillin resistance gene. Constructs from Twist in the pT7rrnB backbone were transformed into chemically competent *E. coli* Dh10β cells, grown in 50-mL cultures in the presence of 100 μg/mL Carbenicillin. Plasmids were then purified using the ZymoPure II Plasmid Miniprep Kit. The resulting plasmids were further purified via ethanol precipitation, using 5 M NH_4_OAc.

Plasmids used for testing of variants in the Squires strain (36, 37) were built using the 7,451-bp pAM552G plasmid. Like the pT7rrnB plasmid, pAM552G carries a copy of the rrnB operon as well as an ampicillin resistance gene. However, in the pAM552 plasmid, rrnB is under control of the phage lambda (pL) promoter, a temperature sensitive promoter. At low temperatures (30 °C), pL is repressed, but it is expressed at higher temperatures (37 °C). Plasmids from Twist were transformed into chemically competent *E. coli* POP2136 cells and miniprepped using the ZymoPURE Plasmid Miniprep Kit. These purified plasmids were then transformed into electrocompetent SQ171fg cells (19, 36). Of note, both plasmids contain an A2058G mutation, which endows the resulting ribosome with Erythromycin (Ery) resistance. Ery is used for the *in vivo* selection.

### Strain culture & harvest

S150 lysates, total protein of the 70S ribosome (TP70) and T7 RNA Polymerase were prepared as previously reported (22, 34). 10 mL of an overnight culture of *E. coli* (MRE600 strain) cells were added into a liter of 2xYTPG medium (2xYTP with 18 g/L of glucose) and grown at 37 °C with shaking at 250 rpm until OD600 reached 3. Culture was spun down at 5,000xg for 10 minutes and kept on ice between all transfer steps. Supernatant was removed and pellet was resuspended in S30 buffer (10 mM TrisOAc pH=8.2, 14 mM Mg(OAc)_2_, 60 mM KOAc). Solution was spun at 10,000xg for 3 minutes twice more, removing supernatant between each spin and resuspending in 40 mL of fresh S30 buffer. After the third spin, pelleted were weighed and flash frozen with liquid nitrogen before storing at -80 °C. S30 Buffer was then added at a ratio of 5 mL per 1g of cell mass, and cells resuspended by vortexing until fully thawed. 100 μL of HALT Protease Inhibitor Cocktail was added per 10 mL cell suspension, and 75 μL of Takada Recombinant RNase Inhibitor was added per 4 grams of dry cell mass. Cells were lysed at ∼25,000 psi with a C3 Avestin Homogenizer and a second aliquot of Takada Recombinant RNase Inhibitor was added at a ratio of 75 μL per 4 grams of initial pellet. Homogenized cells were pelleted at 30,000xg at 4 °C for 30 minutes. Supernatant was recovered and spun a second time at 30,000xg at 4 °C for 30 minutes. Supernatant (S30 extract) was recovered for S150 extract preparation and layered on top of an equivalent volume of sucrose cushion buffer (20 mM Tris– HCl (pH 7.2 at 4 °C), 100 mM NH_4_Cl, 10 mM MgCl_2_, 0.5 mM EDTA, 2 mM DTT, 37.7% sucrose) in Ti70 tubes. Samples were then ultracentrifuged at 90,000xg for 18 hours, after which the supernatant was transferred into fresh Ti70 tubes and spun at 150,000xg for 3 hours and pellets were gently washed with buffer C (10 mM Tris–OAc (pH 7.5 at 4 °C), 60 mM NH_4_Cl, 7.5 mM Mg(Oac)_2_, 0.5 mM EDTA, 2 mM DTT). Ribosome concentration in the pellets was measured using A_260_ Nanodrop measurements (1 *A*_260_ unit of 70S = 24 pmol 70S). After the second spin, the top 2/3 of the supernatant was collected and transferred into MWCO=3.5 K dialysis tubing (SnakeSkin) and dialyzed 2 × 1.5 hours x 3 L of S150 Extract Buffer at 4 °C. For the 3^rd^ dialysis, 3L of fresh S150 Extract Buffer was used to dialyze overnight (12-15 hours). S150 extract was concentrated using Centripreps (3 kDa MWCO) until A_260_=25 and A_280_=15. Extract was aliquoted and flash frozen in liquid nitrogen. TP70 was prepared as previously described (23).

### iSAT reactions

5 μL iSAT reactions were performed in 384-well nunc_267461 plates, with 4 replicates per reaction, and set up as previously described (12, 22, 34). The Echo 525 Acoustic Liquid Handler was used to aliquot reaction components into the wells. Reaction components were prepared in two separate mixtures: 1) the DNA plasmid with a small amount of premix to enable better liquid handling, and 2) the remaining reaction components. Premix was mixed with DNA at a volume ratio of 1:2.2 uL premix:DNA so to enable consistent results by increasing viscosity. Reagent mix 2, containing the S150 extract, was added into the wells initially, and then the DNA plasmid mix was aliquoted into each well. Reactions were run in a plate reader at 37 °C, reading absorbance (Excitation: 485 nm, Emission: 528 nm) every 15 minutes and with constant shaking for 15 hours. 40% PEG8000 (Sigma-Aldrich P1458-23ML) was added into the reaction premix for a final volume of 10%; 1M DTT was added at a final volume of 0.2%.

### Squires transformations and plasmid selections

Electrocompetent Squires cells (SQ171fg) cells containing a pCSacB-KanR plasmid (19, 37) were prepared and stored in 50 μL aliquots. The Squires strain is a modified *E. Coli* strain that has all seven ribosomal operons deleted from the genome. Instead, a ribosomal operon, rrnB, exists on a plasmid that also carries a selection marker. When plasmids carrying the mutated ribosomal operon of interest are transformed into the cell, the original SacB-rrnB plasmid can be removed. 50 μg of purified pAM552G plasmids was transformed into 50 μL of cells. Cells were recovered in 850 μL of SOC in a 1.5-mL microcentrifuge tube at 37 °C for 1 hour, while shaking at 250 rpm. After 1 hour, 270 μL of the cell recovery was added to 2 mL of Super Optimal broth with Catabolite repression (SOC) containing 50 μg/mL Carbenicillin (Cb) and 0.25% sucrose in a 14-mL plastic culture tube. Tubes were incubated at 37 °C overnight, for 16-18 hours. The tubes were then spun down at room temperature for 5 minutes at 4,000xg. 2 mL of clear supernatant was removed, leaving the cell pellet to be concentrated into the remaining 300 μL. This concentrated cell suspension was plated on LB plates containing 5% sucrose, 50 μg/mL Cb, and 20 μL/mL Ery. Plates were incubated at 37 °C until colonies appeared. Once colonies appeared, 8 colonies were picked from each plate and spotted onto two plates, one LB-Cb_50_ plate, and one LB-Kan_50_ plate. Colonies that grew successfully on the Cb_50_ plate but not on the Kan_50_ plate were picked and grown up overnight to be midiprepped using the ZymoPURE II Plasmid Midiprep Kit. Midiprepped plasmids were sequenced to confirm that the 23S sequences were as expected. Constructs that did not yield colonies on the Lb-Suc_5%_-Cb_50_-Ery_20_ plates were transformed two subsequent times to ensure that the construct did not support life. Constructs that did not yield “clean” colonies were troubleshot by picking and spot plating more colonies, and if this was done again without success, transformations were attempted a total of three times before concluding that the construct was not able to support life *in vivo*.

### Spot growth experiment

SQ171fg strains containing the mutated ribosomes of interest were grown overnight in 3-mL cultures, with Cb_50_. In the morning, the OD_600_ of each culture was measured, and normalized to an OD_600_ of 1. Four ten-fold dilutions of each construct were prepared (OD=0.1, 0.01, 0.001, 0.0001). 3 μL of each dilution was carefully pipetted onto a Cb_50_ plate. Plates were incubated at 30 °C and 37 °C and imaged as soon as a construct at the most dilute concentration showed cell growth. Spot growth experiments were completed three separate times to ensure consistent results.

### Cloning and selection of randomized H75 library

Primers were designed with randomized nucleotides at the Helix 75 library location. Two PCRs were performed using primers with nucleotides randomized at the correct location (**Supplementary Table S1**). These two fragments were ligated using Gibson assembly and transformed into chemically competent Dh10B cells. The transformation was allowed to recover for one hour at 37 °C before being plated on Cb50 and grown overnight at 37 °C. Fourteen colonies were picked randomly, and plasmids purified as reported above to test in iSAT.

## Results

Our goal was to establish a high-throughput, computationally guided approach to identify functionally active mutant ribosomes. As model regions to mutate, we focused on important helices within the PTC, as the PTC plays the central role in the dynamic process of peptide bond formation. Specifically, we selected Helix 75 (H75), Helix 73 (H73), Helix 91 (H91), and Helix 92 (H92), and combinations thereof. H75 is known to play a role in the assembly of the nascent polypeptide exit tunnel (5) which is essential to proper polypeptide folding. Along with helices 76 and 79, H75 forms the base of the L1 stalk (38), which facilitates binding, movement, and release of deacylated tRNAs (39). H73 is in the aminoacyl site (A-site) of the PTC and makes contacts with ribosomal (r-)protein L3 (40, 41). This RNA-protein interaction is the closest approach of any amino acid into the PTC (40). H91 and H92 together form one side of the “accommodation corridor,” where aminoacylated tRNAs are directed into the PTC in a specific orientation. H73 and H91 are also interesting candidates for mutation as they share a protein interaction with r-proteins L19 and L14 which could potentially serve as links between the two helices and drive epistatic interactions (42).

### Establishing Stepwise Monte Carlo library selection on Helix 75

We first established the ability of SWM to computationally design and select mutations in H75 of the 23S rRNA, which is at the edge of the PTC. In short, SWM searches libraries by using an add-and-delete move method with stochastic sampling and produces a Rosetta all-atom energy score, which is a linear combination of scaled statistical and physics-based energy terms (31, 43). This optimized energy score serves as a metric to understand and compare sequence stabilities. These simulations and resulting scores account for interactions with nearby residues, whether RNA, protein, ion, or water. For H75, we created a library of rRNA variants by selecting eight nucleotides that make up the center of the helix and permitted each residue to be anything other than its identity in the wildtype (WT) ribosome (**Figure 1A**). Using this problem definition, we ran 10,000 SWM design simulations and selected 50 resulting sequences whose scores spanned the energy score range to build (**Supplementary Table S1**).

**Figure 1.**
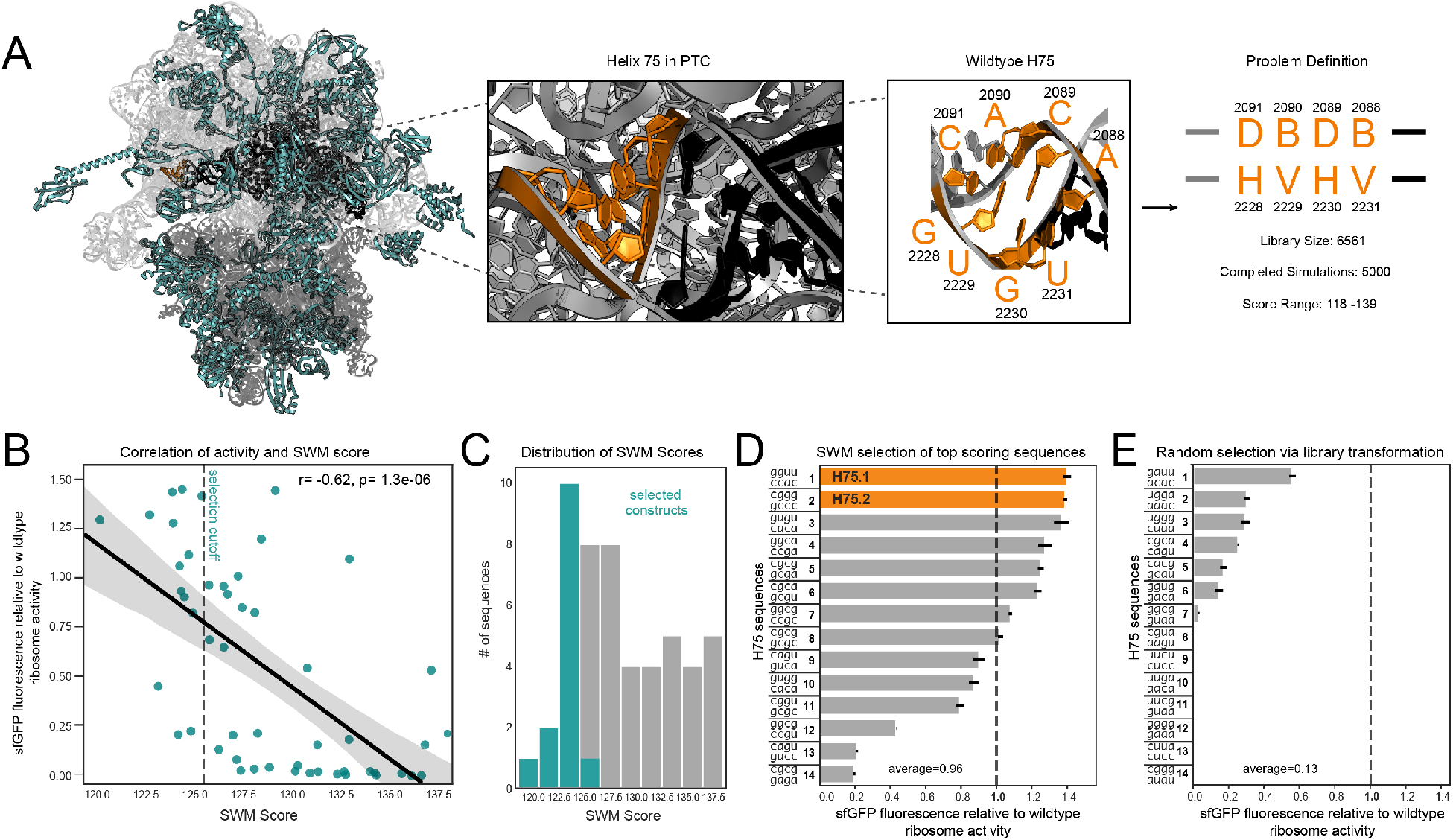
Application of SWM to selection of H75 variants yields high-performing mutants in iSAT. (**A**) Structure and library design for H75. H75, highlighted in orange, sits on the edge of the PTC, which is highlighted in black. Ribosomal proteins are highlighted in teal. (**B**) Correlation between iSAT activity of mutants and their SWM scores. Dashed line indicates SWM score cutoff for selection. (**C**) Selection scheme for constructs based on their SWM scores. (**D**) iSAT activities of SWM selected constructs. (**E**) iSAT activities of randomly selected mutants. Dashed line highlights wild type activity. Data are presented of means of n = 3 experiments with standard deviation shown.

With these SWM-scored mutant rRNA sequences at hand, we next tested their activity in the high-throughput iSAT platform. iSAT co-activates the processes of rRNA synthesis and processing, ribosome assembly, and translation in a one-pot *in vitro* reaction (22). Translational activity was quantified by monitoring superfolder green fluorescent protein (sfGFP) expression. The 50 rRNA mutant sequences were tested in iSAT to see if the SWM conformation score was correlated to iSAT performance. Maximum sfGFP synthesized after a 16-hour iSAT reaction incubated at 37 °C was measured for each construct and normalized relative to that of the wildtype ribosome control. We observed a correlation (r=0.62) between performance in iSAT and SWM score (**Figure 1B**). Lower scores, which indicate a more stable rRNA structural conformation, were more likely to yield active ribosomes, a result consistent with our recent work (44). Given this correlation, we moved forward with using SWM score as a metric for selection of highly active ribosomes.

While we initially tested constructs with a broad range of SWM scores to explore the correlation between score and activity, we wanted to confirm that using this relationship as a selection criterion would enable identification of highly active constructs. We therefore picked sequences with scores in the top 30% (**Figure 1C, Figure 1D**) and selected an equal number of constructs from a randomized library as a negative control using randomized primers (**Figure 1E, Supplementary Table S2**) to test in iSAT. Of the constructs selected using our scoring metric, 7/14 of the selected sequences outperformed the WT sequence in iSAT. The average relative activity of SWM selected sequences was 0.96 (**Figure 1D**) compared to the average of 0.13 for sequences randomly selected from the negative control library (**Figure 1E**). This indicates that our selection method allows for the identification of highly active variants in iSAT and highlights the flexibility of the PTC to mutations when using the SWM design strategy. Additionally, many of the selected sequences were non-trivial solutions, in that they did not maintain the base pairing pattern of the WT H75. For example, H75.1 (cggu,gcgc), which can have only two Watson-Crick (WC) base pairing interactions as opposed to the four WC base pairs in the WT helix, shows near WT activity in iSAT. We also see that many of the selected sequences, such as H75.1, H75.4, and H75.5, do not have a WC interaction at the fourth position between residues 2228 and 2091. Although some crystal structures have found this pair to be closely interacting (45), other studies, specifically mapping secondary structure of the 23S rRNA using base-pairing and stacking interactions (46) show that G2228 is pulled away from C2091 due to a bulging motif at the base of Helix 79. This indicates that our computational modeling approach was able to detect and account for the additional flexibility of this base pair and score sequences that left this region less rigid favorably.

### Application of SWM to select for active mutated helices in the A and P sites

We next sought to use SWM design and selection on additional motifs within the PTC to assess mutational flexibility and find novel ribosomal mutants (**Figure 2A**). We chose H73, H91, and H92, three helices that are known to play roles of varying importance to the ribosome’s dynamic activity. Using a similar approach as for H75, we ran design simulations for H73, H91 and 92, and as above selected 14 sequences from each library that had energy scores in the best 30% of scores to build and test in iSAT (**Supplementary Table S3, Supplementary Table S4**). All three simulations provided us with viable 23S rRNA sequences, and multiple sequences outperformed the WT control in iSAT (**Figure 2B-D**). The observed trends highlighted that the ribosomal mutants do not have to exactly mimic WT base pairing patterns in order to be highly active. For example, variants H73.3, H73.4 and H73.5 (**Figure 2B**), which are all at least as active as WT in iSAT, have at least one base pair that is not a canonical WC base pair. Additionally, our data supported previous findings that ribosome activity is highly sensitive to even small sequence changes. The identity of even one non-WC base pair can significantly affect activity; H91.3 and H91.4 are nearly identical, but H91.4 is significantly less active because of a single nucleotide change that converts a c:c pair to an a:c pair (**Figure 2C**). This suggests that 2539C is more favorable than 2539A, perhaps due to interactions occurring between 2539 and nearby rRNA or r-protein residues; or the result of slightly stronger hydrogen bonding of a C*C bond than an A*C bond (47). Surprisingly, between H91.3 and H91.6, we observed that changing the first base pair from a:u to c:g and the fourth from c:c to c:g actually leads to an almost 2-fold knockdown of activity. The selected sequences from H92 were the least active, indicating that the ribosome is more sensitive to changes in this helix (**Figure 2D**). This is likely because H92 is specifically recognized by DbpA, an RNA helicase that is known to play an important role in ribosome assembly (48, 49). In addition, we find that perfect base pairing does not guarantee high performance; H92.4 has perfect base pairing but less than a third of the iSAT activity of H92.1. A library design for H92 that includes additional randomized residues could allow for better compensatory mutations to be identified in this region.

**Figure 2.**
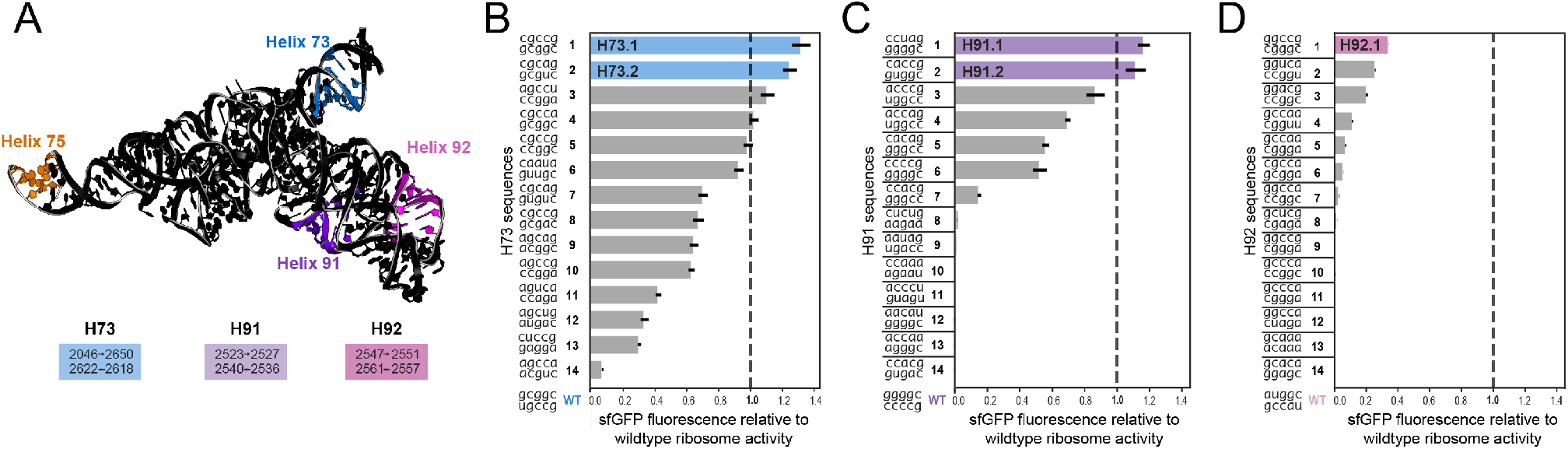
Design of additional helices using SWM yields multiple highly active mutants. (**A**) Highlighted locations and residue numbers of additional helices within the PTC. (**B**) iSAT results for selected H73 variants. (**C**) iSAT results for selected H91 variants. (**D**) iSAT results for selected H92 variants. Dashed line indicates the activity of WT, normalized to 1. Dashed line highlights wild type activity. Data are presented of means of n = 3 experiments with standard deviation shown.

### Combinatorial analysis of top performing sequences highlights complex epistatic interactions in the PTC

We next wondered if combining mutations across different helices in the PTC would lead to compensatory, beneficial phenotypes. To test this, we selected top-performing sequences from each library (highlighted in **Figure 1D** and **Figure 2B-D**) and constructed all possible combinations of the four library region sequences including WT, yielding a total of 54 combinatorial rRNA constructs to test in iSAT. While none of the constructs with all four library locations mutated produced detectable concentrations of sfGFP, many of the 3-way combinations were highly functional and even competitive iSAT activity with the WT control (**Figure 3**; **Supplementary Table S5, Supplementary Figure S1**).

**Figure 3.**
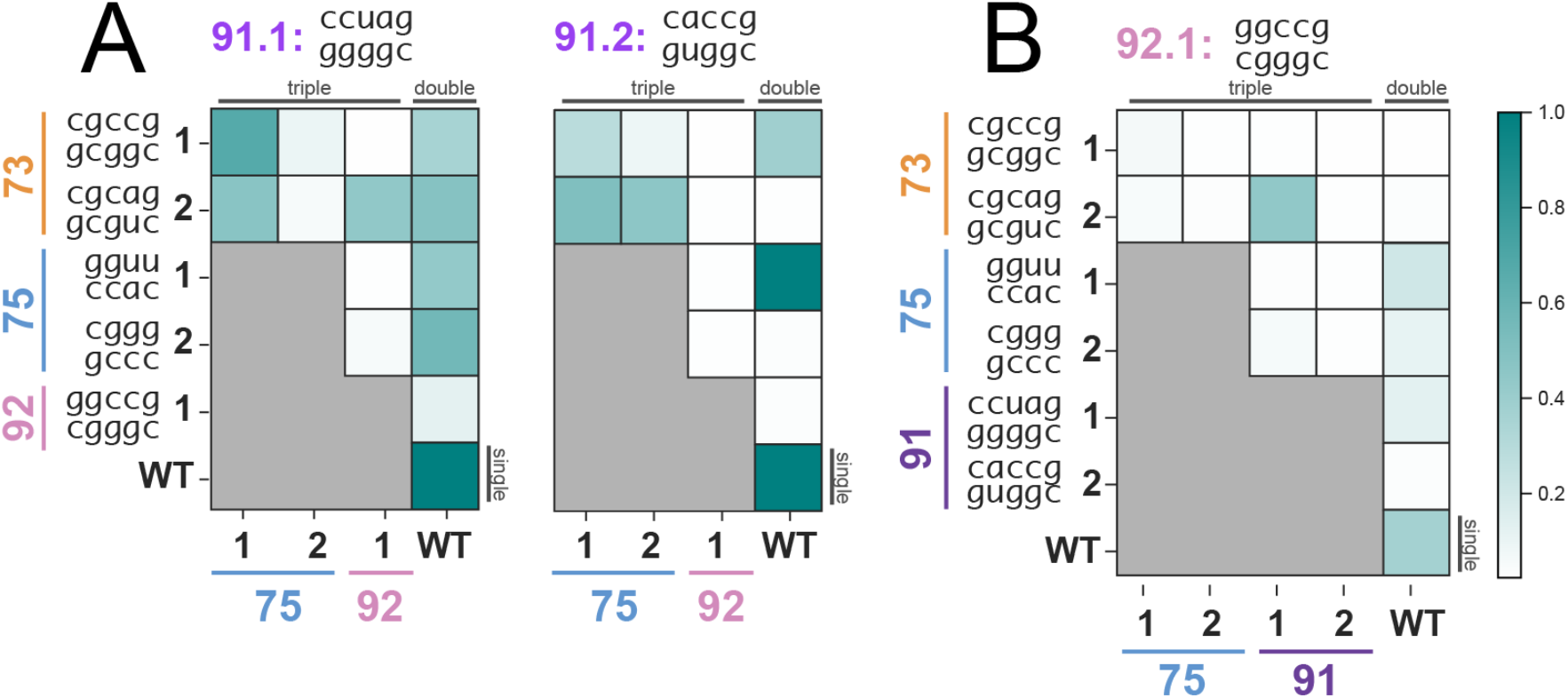
iSAT activities of two and three-way PTC mutant combinations yield constructs with varying activities and highlight epistatic interactions. (**A**) H91 mutations kept constant; all constructs shown include a mutated H91, as indicated in title. (**B**) H92.1 mutant included in all combinations. Data represent the average of n = 3 independent experiments.

The results uncovered interesting epistatic interactions between helices that highlight the highly interacting structure and complexity of the PTC. iSAT activity could be recovered by adding additional mutated helices in some cases. For instance, we observed that H91.2 combined with H73.2 is inactive, but when further combined with either H75.1 or H75.2 the activity relative to WT increases to 0.50 (**Figure 3A, Supplementary Table S5**). Although H73/H91 and H75 sit at opposite ends of the PTC, the addition of 8 mutated residues at H75 unexpectedly complements mutated H91.2 and H73.2, rendering them compatible. In another example, we showed that the double mutant of H91.2 combined with H73.1 is moderately active while H91.2 with H73.2, which has only one base pair difference from H73.1, has no detectable activity. H73’s effects on H91 are different when looking at the H91.1 mutant; H91.1 has 0.50 relative activity with both H73.1 and H73.2. The sensitivity of the H91 variants to small changes in H73 sequence may be due to shared interactions with proteins L14 and L19, which are thought to play important roles in particle assembly (42). Similarly, while H91.1 is active as a double mutant with H75.1 or H75.2, H91.2 has very high activity when paired with H75.1 but none with H75-- even though H75.1 and H75.2 have almost identical activities individually (**Figure 1D, Supplementary Table S3**). We were surprised to see that all triple mutants including H92.1 are near inactive except for its combination with H73.2 and H91.1 (**Figure 3B**). In fact, the addition of the 20 mutations from H73.2 and H91.1 endows the ribosome with significantly higher activity than in any double or single mutant of those regions, indicating a sensitive relationship between the three helices. This could be explained by role of the L3 protein, which acts as a dynamic switch to coordinate binding of elongation factors and has been reported to interact with helices 73, 91 and 92 as amino-acid charged tRNAs are introduced into the A-site and shuttled to the P-site (40). These mutant combinations highlight four key findings: (i) there exist interesting, previously unexplored relationships between helices in the PTC, (ii) considering dynamic and distal interactions in rRNA is essential for successful rRNA library design, (iii) ribosome activity is sensitive to even single base pair differences in helices, and (iv) the PTC is highly flexible to large-scale mutations despite its sequence conservation.

### Highly mutated, computationally designed ribosomes support life in vivo

We then transformed all combinatorial constructs and single mutants into *E. coli* to test whether these mutant ribosomes could support translation of the *E. coli* proteome. We used a previously described selection scheme (12, 37). In short, plasmids conferring carbenicillin resistance and encoding mutant ribosomes were transformed into the *E. coli* SQ171fg strain (50), which lacks chromosomal rRNA alleles and lives on the pCSacB plasmid carrying a WT rRNA operon. The pCSacB plasmid also contains a counter-selectable marker sacB gene which confers sucrose sensitivity and a kanamycin resistance cassette. Thus, by growing transformed SQ171fg cells in the presence of carbenicillin and sucrose and assessing kanamycin sensitivity, individual colonies can be selected that support life from mutant ribosomes.

While most combinatorial constructs were not able to support life, many of them were successful and closely resembled WT phenotypically (**Figure 4A**). We found that strain C22, containing H73.1 and H91.1, grew slowly, but when combined with H75.1 (strain C33), growth was improved significantly, matching the phenotype observed *in vitro* (**Figure 3A**). Likewise, H91.1 alone grew more slowly than its combination with either H75.1 (C55) or H75.2 (C60). Of the constructs that had greater than ∼1/3 relative activity to WT in iSAT, more than 80% were able to support life; thus, our data suggest that iSAT activity can inform which ribosome mutants might support life. However, there were some exceptions. For example, the H73.1-H75.1-H91.1 combination was highly active in iSAT but did not support life. Conversely, strain C39 was able to support cell growth despite its low activity in iSAT. These disparities are likely due to differences in the concentrations of the many dozens of assembly factors involved in ribosome biogenesis in either environment (51). We also measured the growth profiles of these strains at a lower temperature (30 °C) to observe any phenotypic changes that may be more pronounced in suboptimal growth conditions (**Figure 4B**). Surprisingly, strains C33 and C39, which showed slightly impaired growth at 37 °C, grew significantly more robustly than WT at the lower temperature. This suggests that the mutations in strains C33 and C39 may lead to improved folding and assembly in a low temperature cell environment. Strains C16, C33 and C39 all harbor ribosomes that have 10% of their PTCs mutated from WT and are still able to support cell survival and growth.

**Figure 4.**
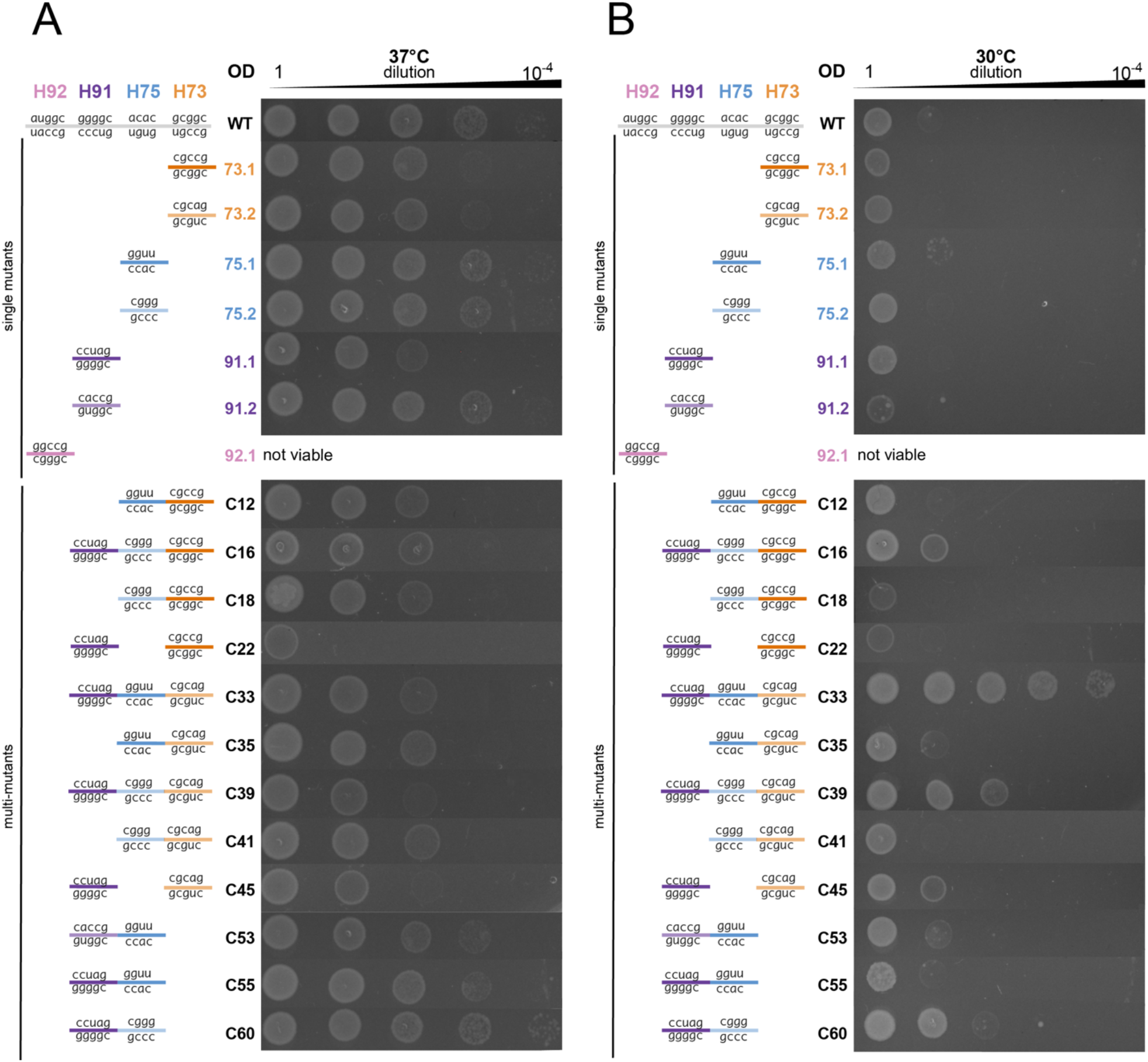
Spot growth assays of controls and combination constructs show that many mutants can support cell growth competitive with wildtype. (**A**) Spot growth assay at 37 °C (**B**) Spot growth assay at 30 °C. Data representative of n=3 independent experiments.

## Discussion

Here, we developed a computational rRNA structure prediction method to select for highly active ribosomal PTC mutants from complex libraries and explore previously unidentified epistatic interactions between distal helices in the PTC. Our work has several key features. First, we showed that we can use SWM to successfully select for highly active mutants using an all-atom energy score, and that this approach can serve as a tool to design mutants that not only do not disrupt ribosome function, but even improve it. This finding is consistent with current understanding of the importance of rapid rRNA folding for ribosome assembly and function (52), which has long been understood to be a function of the molecule’s minimum free energy (53, 54). By combining computationally designed helix mutants, we built functional ribosomes with up to 30 mutations, the most highly mutated designed PTCs using SWM to our knowledge, showing that the PTC is much more flexible to mutation than previously thought. Many of these multi-mutants were able to support life; some strains showed improved growth phenotypes to that of a strain carrying a WT ribosome. Additionally, we observed that constructs that were more active *in vitro* had a higher probability of being able to support life in cells and identified a general rule for predicting *in vivo* success as a function of iSAT performance, which appears to be agnostic to helix location.

Second, our approach enabled library design through the lens of three-dimensional structure. We believe this feature is important for rRNA design, as nucleotides that are distant in sequence space are often highly interacting in three-dimensional space and mutating a single residue can have off-target effects on other rRNA motifs. SWM also allows for unbiased library assessment, which is experimentally challenging due to inherent biases of primer synthesis and template:primer interactions. By computationally exploring large libraries of mutations in the PTC computationally, we allowed the ribosome to explore the folding energy landscape and find alternative minima that retained—and sometimes improved—function.

Third, select combinatorial mutants highlighted previously unidentified epistatic interactions between helices in the PTC. For example, activities of A-site helix mutants (H91, H92, H73) were strongly affected by mutations in the E-site (H75). This effect was highly sensitive to even small sequence changes; a single base pair change in H73, for example, rendered the multi-mutant with H91.2/H75.2 incapable of producing sfGFP. While H73 and H91 have been reported to be related because of their proximity within the A-site and interactions with the dynamic r-protein L3 as well as proteins L14 and L19 (40, 42), their relationship with H75 has not been documented. This finding suggests that these rRNA motifs are co-dependent, perhaps due to the activities of additional r-proteins working together dynamically, and that their mutations can render the ribosome inactive if incompatible. Notably, combining highly active mutants does not guarantee that the resulting multi-mutant will be active; as seen with H75.2 and H91.1, which individually both outperformed the WT ribosome in iSAT, their combination abolished iSAT activity (**Figure 2B**). This finding emphasizes the high interconnectivity of the PTC and the need to approach engineering it through a wide lens; identifying the most active small-scale mutants individually to later combine is an oversimplification of the design challenge. Thus, high-throughput screens will likely be required to test diverse mutations in combination to enable discovery of the most promising multi-mutants. The unexpected relationships between distant helices of the PTC underscore the dynamic activity of the ribosomal active site and help improve our understanding of how to account for these kinds of interactions in future ribosome engineering efforts.

Looking forward, we anticipate that energy-based structure predictions such as SWM will be important to facilitate ribosome design. This holds promise to advance our understanding of rRNA function and molecular translation, as well as accelerate efforts in making modified ribosomes with expanded functions for chemical and synthetic biology.

## Data Availability

Methods and input files used to run SWM simulations are available at https://github.com/everyday847/ptc_swm_modeling. All other data is available in the main text or in the Supplementary Information.

## Acknowledgements

The authors would like to thank Dr. Gyorgy Bagnigg and Argonne National Labs for access to the advanced computing center. The authors are also thankful to Ramya Rangan and Phillip Pham for help beta testing the computational pipeline. C.K. was supported by the National Science Foundation Graduate Research Fellowship Program under grant no. DGE-1842165. This work was also supported by the Army Research Office (W911NF-16-1-0372) and Army Contracting Command (W52P1J-21-9-3023). The U.S. Government is authorized to reproduce and distribute reprints for Governmental purposes notwithstanding any copyright notation thereon. The views and conclusions contained herein are those of the authors and should not be interpreted as necessarily representing the official policies or endorsements, either expressed or implied, of the U.S. Government.

## Author Contributions

C.K., A.M.W., D.S.K., R.D., and M.C.J. conceived the project idea and planned experiments. A.M.W. and R.D. wrote the stepwise Monte Carlo design code in Rosetta. C.K. and A.M.W. ran computational simulations. C.K. and A.C.W. conducted experiments. C.K. and A.K. analyzed the data and made figures. C.K. wrote the paper. A.K., M.C.J., A.M.W., D.S.K., A.C.W. and R.D. revised and edited the paper.

## References

1. Isenbarger, T.A., Carr, C.E., Stewart Johnson, S., Finney, M., Church, G.M., Gilbert, W., Zuber, M.T., Ruvkun, G., Carr, C.E., Johnson, S.S., et al. (2008) The Most Conserved Genome Segments for Life Detection on Earth and Other Planets. 10.1007/s11084-008-9148-z.

2. Nissen, P., Hansen, J., Ban, N., Moore, P.B. and Steitz, T.A. (2000) The structural basis of ribosome activity in peptide bond synthesis. Science, 289, 920–930.

3. Steitz, T.A. and Moore, P.B. (2003) RNA, the first macromolecular catalyst: the ribosome is a ribozyme. Trends Biochem. Sci., 28, 411–418.

4. Cech, T.R. (2000) The Ribosome Is a Ribozyme. Science (80-.)., 289, 878–879.

5. Wilson, D.M., Li, Y., LaPeruta, A., Gamalinda, M., Gao, N. and Woolford, J.L. (2020) Structural insights into assembly of the ribosomal nascent polypeptide exit tunnel. Nat. Commun. 2020 111, 11, 1–15.

6. Kaczanowska, M. and Rydén-Aulin, M. (2007) Ribosome Biogenesis and the Translation Process in Escherichia coli. Microbiol. Mol. Biol. Rev., 71, 477–494.

7. Cochella, L. and Green, R. (2004) Isolation of antibiotic resistance mutations in the rRNA by using an in vitro selection system. Proc. Natl. Acad. Sci. U. S. A., 101, 3786–3791.

8. Dedkova, L.M., Fahmi, N.E., Golovine, S.Y. and Hecht, S.M. (2006) Construction of modified ribosomes for incorporation of D-amino acids into proteins. Biochemistry, 45, 15541– 15551.

9. Dedkova, L.M., Fahmi, N.E., Paul, R., Del Rosario, M., Zhang, L., Chen, S., Feder, G. and Hecht, S.M. (2012) β-puromycin selection of modified ribosomes for in vitro incorporation of β-amino acids. Biochemistry, 51, 401–415.

10. D’Aquino, A.E., Kim, D.S. and Jewett, M.C. (2018) Engineered ribosomes for basic science and synthetic biology. Annu. Rev. Chem. Biomol. Eng., 9, 311–340.

11. Thompson, J., Kim, D.F., O’Connor, M., Lieberman, K.R., Bayfield, M.A., Gregory, S.T., Green, R., Noller, H.F. and Dahlberg, A.E. (2001) Analysis of mutations at residues A2451 and G2447 of 23S rRNA in the peptidyltransferase active site of the 50S ribosomal subunit. Proc. Natl. Acad. Sci., 98, 9002–9007.

12. D’Aquino, A.E., Azim, T., Aleksashin, N.A., Hockenberry, A.J., Krüger, A. and Jewett, M.C. (2020) Mutational characterization and mapping of the 70S ribosome active site. Nucleic Acids Res., 48, 2777–2789.

13. Rakauskaite, R. and Dinman, J.D. (2011) Mutations of highly conserved bases in the peptidyltransferase center induce compensatory rearrangements in yeast ribosomes. Rna, 17, 855–864.

14. Green, R., Samaha, R.R. and Noller, H.F. (1997) Mutations at nucleotides G2251 and U2585 of 23 S rRNA perturb the peptidyl transferase center of the ribosome. J. Mol. Biol., 266, 40–50.

15. Long, K.S., Munck, C., Andersen, T.M.B., Schaub, M.A., Hobbie, S.N., Böttger, E.C. and Vester, B. (2010) Mutations in 23S rRNA at the peptidyl transferase center and their relationship to linezolid binding and cross-resistance. Antimicrob. Agents Chemother., 54, 4705–4713.

16. Dutheil, J.Y., Jossinet, F. and Westhof, E. (2010) Base pairing constraints drive structural epistasis in ribosomal RNA sequences. Mol. Biol. Evol., 27, 1868–1876.

17. Fried, S.D., Schmied, W.H., Uttamapinant, C. and Chin, J.W. (2015) Ribosome Subunit Stapling for Orthogonal Translation in E.coli. Angew. Chem. Int. Ed. Engl., 54, 12791.

18. Aleksashin, N.A., Szal, T., d’Aquino, A.E., Jewett, M.C., Vázquez-Laslop, N. and Mankin, A.S. (2020) A fully orthogonal system for protein synthesis in bacterial cells. Nat. Commun., 10.1038/s41467-020-15756-1.

19. Orelle, C., Carlson, E.D., Szal, T., Florin, T., Jewett, M.C. and Mankin, A.S. Protein synthesis by ribosomes with tethered subunits. 10.1038/nature14862.

20. Kolber, N.S., Fattal, R., Bratulic, S., Carver, G.D. and Badran, A.H. Orthogonal translation enables heterologous ribosome engineering in E. coli. 10.1038/s41467-020-20759-z.

21. Youngman, E.M. and Green, R. (2005) Affinity purification of in vivo-assembled ribosomes for in vitro biochemical analysis. Methods, 36, 305–312.

22. Jewett, M.C., Fritz, B.R., Timmerman, L.E. and Church, G.M. (2013) In vitro integration of ribosomal RNA synthesis, ribosome assembly, and translation. Mol. Syst. Biol., 9, 678.

23. Hammerling, M.J., Fritz, B.R., Yoesep, D.J., Kim, D.S., Carlson, E.D. and Jewett, M.C. (2020) In vitro ribosome synthesis and evolution through ribosome display. Nat. Commun., 11.

24. Huang, S., Aleksashin, N.A., Loveland, A.B., Klepacki, D., Reier, K., Kefi, A., Szal, T., Remme, J., Jaeger, L., Vázquez-Laslop, N., et al. (2020) Ribosome engineering reveals the importance of 5S rRNA autonomy for ribosome assembly. Nat. Commun., 11, 1–13.

25. Yassin, A. and Mankin, A.S. (2007) Potential New Antibiotic Sites in the Ribosome Revealed by Deleterious Mutations in RNA of the Large Ribosomal Subunit *. 10.1074/jbc.M703106200.

26. Neylon, C. (2004) Chemical and biochemical strategies for the randomization of protein encoding DNA sequences: library construction methods for directed evolution. Nucleic Acids Res., 32, 1448.

27. Kim, D.S., Watkins, A., Bidstrup, E., Lee, J., Topkar, V., Kofman, C., Schwarz, K.J., Liu, Y., Pintilie, G., Roney, E., et al. (2022) 3D-structure-guided evolution of a ribosome with tethered. Nat. Chem. Biol.

28. Sato, N.S., Hirabayashi, N., Agmon, I., Yonath, A. and Suzuki, T. (2006) Comprehensive genetic selection revealed essential bases in the peptidyl-transferase center. Proc. Natl. Acad. Sci. U. S. A., 103, 15386–15391.

29. Dedkova, L.M. and Hecht, S.M. (2019) Expanding the Scope of Protein Synthesis Using Modified Ribosomes. J. Am. Chem. Soc., 141, 6430–6447.

30. Dedkova, L.M., Fahmi, N.E., Paul, R., Del Rosario, M., Zhang, L., Chen, S., Feder, G. and Hecht, S.M. (2012) B-Puromycin Selection of Modified Ribosomes for in Vitro Incorporation of B-Amino Acids. Biochemistry, 51, 401–415.

31. Watkins, A.M., Geniesse, C., Kladwang, W., Zakrevsky, P., Jaeger, L. and Das, R. (2018) Blind prediction of noncanonical RNA structure at atomic accuracy. Sci. Adv., 4, 1–13.

32. Boniecki, M.J., Lach, G., Dawson, W.K., Tomala, K., Lukasz, P., Soltysinski, T., Rother, K.M. and Bujnicki, J.M. (2016) SimRNA: a coarse-grained method for RNA folding simulations and 3D structure prediction. Nucleic Acids Res., 44, e63.

33. Liu, Y., Fritz, B.R., Anderson, M.J., Schoborg, J.A. and Jewett, M.C. (2015) Characterizing and alleviating substrate limitations for improved in vitro ribosome construction. ACS Synth. Biol., 4, 454–462.

34. Fritz, B.R. and Jewett, M.C. (2014) The impact of transcriptional tuning on in vitro integrated rRNA transcription and ribosome construction. Nucleic Acids Res., 42, 6774–6785.

35. Fritz, B.R., Jamil, O.K. and Jewett, M.C. (2015) Implications of macromolecular crowding and reducing conditions for in vitro ribosome construction. Nucleic Acids Res., 43, 4774–4784.

36. Quan, S., Skovgaard, O., McLaughlin, R.E., Buurman, E.T. and Squires, C.L. (2015) Markerless Escherichia coli rrn deletion strains for genetic determination of ribosomal binding sites. G3 Genes, Genomes, Genet., 5, 2555–2557.

37. Asai, T., Condon, C., Voulgaris, J., Zaporojets, D., Shen, B., Al-Omar, M., Squires, C. and Squires, C.L. (1999) Construction and initial characterization of Escherichia coli strains with few or no intact chromosomal rRNA operons. J. Bacteriol., 181, 3803–3809.

38. Réblová, K., Šponer, J. and Lankaš, F. (2012) Structure and mechanical properties of the ribosomal L1 stalk three-way junction. Nucleic Acids Res., 40, 6290–6303.

39. Cornish, P. V., Ermolenko, D.N., Staple, D.W., Hoang, L., Hickerson, R.P., Noller, H.F. and Ha, T. (2009) Following movement of the L1 stalk between three functional states in single ribosomes. Proc. Natl. Acad. Sci., 106, 2571–2576.

40. Meskauskas, A. and Dinman, J.D. (2015) Ribosomal Protein L3: Gatekeeper to the A Site. Mol. Cell, 10.1016/j.molcel.2007.02.015.

41. Meskauskas, A. and Dinman, J.D. (2008) Ribosomal protein L3 functions as a ‘rocker switch’ to aid in coordinating of large subunit-associated functions in eukaryotes and Archaea. Nucleic Acids Res., 36, 6175–6186.

42. Davies, C., White, S.W. and Ramakrishnan, V. (1996) The crystal structure of ribosomal protein L14 reveals an important organizational component of the translational apparatus. Structure, 4, 55–66.

43. Alford, R.F., Leaver-Fay, A., Jeliazkov, J.R., O’Meara, M.J., DiMaio, F.P., Park, H., Shapovalov, M. V., Renfrew, P.D., Mulligan, V.K., Kappel, K., et al. (2017) The Rosetta All-Atom Energy Function for Macromolecular Modeling and Design. J. Chem. Theory Comput., 13, 3031–3048.

44. Krüger, A., Watkins, A.M., Wellington-Oguri, R., Romano, J., Kofman, C., DeFoe, A., Kim, Y., Anderson-Lee, J., Fisker, E., Townley, J., et al. (2021) Community science designed ribosomes with beneficial phenotypes. bioRxiv, 10.1101/2021.09.05.458952.

45. Noeske, J., Wasserman, M.R., Terry, D.S., Altman, R.B., Blanchard, S.C. and Cate, J.H.D. (2015) High-resolution structure of the Escherichia coli ribosome. Nat. Struct. Mol. Biol., 22, 336–341.

46. Petrov, A.S., Bernier, C.R., Hershkovits, E., Xue, Y., Waterbury, C.C., Hsiao, C., Stepanov, V.G., Gaucher, E.A., Grover, M.A., Harvey, S.C., et al. (2013) Secondary structure and domain architecture of the 23S and 5S rRNAs. Nucleic Acids Res., 41, 7522.

47. Lemieux, S. and Major, F. (2002) RNA canonical and non-canonical base pairing types: a recognition method and complete repertoire. Nucleic Acids Res., 30, 4250.

48. Karginov, F. V. and Uhlenbeck, O.C. (2004) Interaction of Escherichia coli DbpA with 23S rRNA in different functional states of the enzyme. Nucleic Acids Res., 32, 3028.

49. Sharpe Elles,L.M., Sykes, M.T., Williamson, J.R. and Uhlenbeck, O.C. (2009) A dominant negative mutant of the E. coli RNA helicase DbpA blocks assembly of the 50S ribosomal subunit. Nucleic Acids Res., 37, 6503.

50. Carlson, E.D., d’Aquino, A.E., Kim, D.S., Fulk, E.M., Hoang, K., Szal, T., Mankin, A.S. and Jewett, M.C. (2019) Engineered ribosomes with tethered subunits for expanding biological function. Nat. Commun., 10.

51. Williamson, J.R. and Davis, J.H. (2017) Structure and dynamics of bacterial ribosome biogenesis. Philos. Trans. R. Soc. B Biol. Sci., 372.

52. Woodson, S.A. (2008) RNA folding and ribosome assembly. Curr. Opin. Chem. Biol., 12, 667.

53. Chen, S.J. and Dill, K.A. (2000) RNA folding energy landscapes. Proc. Natl. Acad. Sci., 97, 646–651.

54. Doshi, K.J., Cannone, J.J., Cobaugh, C.W. and Gutell, R.R. (2004) Evaluation of the suitability of free-energy minimization using nearest-neighbor energy parameters for RNA secondary structure prediction. BMC Bioinformatics, 5, 1–22.

