## Supplement for "Computationally-guided design and selection of ribosomal active site mutants with high activity"

a. Department of Chemical and Biological Engineering, Northwestern University, Evanston, IL 60208, USA
b. Chemistry of Life Processes Institute, Northwestern University, Evanston, IL 60208, USA
c. Center for Synthetic Biology, Northwestern University, Evanston, IL 60208, USA
d. Department of Biochemistry, Stanford University, Stanford, CA 94305, USA
e. Prescient Design, Genentech, South San Francisco, CA 94080, USA.
f. Department of Physics, Stanford University, Stanford, CA 94305, USA
g. Robert H. Lurie Comprehensive Cancer Center and Simpson Querrey Institute, Northwestern University, Chicago, IL 60611, USA

#### Correspondence

Michael C. Jewett,

### Supplemental Information

#### Supplementary Tables

##### Table S1. iSAT activities, SWM scores, and sequences of constructed H75 mutants.

| Name | Normalized iSAT Activity | Standard Deviation | Top SWM Score | Sequence |
|---|---|---|---|---|
| **H75.1*** | 1.400 | 0.022 | 123.577 | gguu, cacc |
| **H75.2*** | 1.387 | 0.013 | 122.990 | cggg,cccg |
| **H75.3** | 1.367 | 0.041 | 124.748 | gugu,acac |
| **H75.4** | 1.275 | 0.039 | 121.649 | ggca,agcc |
| **H75.5** | 1.250 | 0.017 | 118.696 | cgcg,agcg |
| **H75.6** | 1.234 | 0.021 | 123.039 | cgca,ugcg |
| **H75.7** | 1.079 | 0.008 | 123.966 | ggcg,cgcc |
| **H75.8** | 1.023 | 0.014 | 123.422 | cgcg,cgcg |
| **H75.9** | 0.901 | 0.035 | 123.527 | cagu,acug |
| **H75.10** | 0.871 | 0.026 | 123.707 | gugg,acac |
| **H75.11** | 0.792 | 0.023 | 124.208 | cggu,cgcg |
| **H75.12** | 0.433 | 0.002 | 122.151 | ggcg,ugcc |
| **H75.13** | 0.212 | 0.006 | 124.093 | cagu,ccug |
| **H75.14** | 0.195 | 0.007 | 123.348 | cgcg,agag |
| **H75.15** | 1.394 | 0.028 | 129.105 | guuu,aacc |
| **H75.16** | 1.058 | 0.010 | 133.509 | uaga,uaua |
| **H75.17** | 0.973 | 0.006 | 126.898 | acac,gugu |
| **H75.18** | 0.930 | 0.031 | 125.170 | cggu,cccc |
| **H75.19** | 0.924 | 0.030 | 126.049 | caac,guug |
| **H75.20** | 0.885 | 0.005 | 126.279 | gggu,caca |
| **H75.21** | 0.819 | 0.005 | 127.129 | aggc,acag |
| **H75.22** | 0.794 | 0.024 | 127.892 | cgcg,cccg |
| **H75.23** | 0.661 | 0.013 | 125.199 | cgcg,accg |
| **H75.24** | 0.624 | 0.015 | 126.054 | ggcg,ugac |
| **H75.25** | 0.522 | 0.008 | 131.004 | gugu,caca |
| **H75.26** | 0.511 | 0.007 | 138.373 | auag,caca |
| **H75.27** | 0.201 | 0.006 | 128.066 | cggc,aguc |
| **H75.28** | 0.192 | 0.005 | 126.597 | cagu,cgug |
| **H75.29** | 0.182 | 0.027 | 138.646 | gaag,caca |
| **H75.30** | 0.171 | 0.005 | 133.484 | uacg,ccua |
| **H75.31** | 0.145 | 0.002 | 137.995 | aucg,caca |
| **H75.32** | 0.145 | 0.004 | 131.605 | ggcg,uaac |
| **H75.33** | 0.122 | 0.002 | 125.756 | cggc,acuc |
| **H75.34** | 0.073 | 0.001 | 126.812 | cggu,cgag |
| **H75.35** | 0.043 | 0.003 | 130.445 | gagc,ugca |
| **H75.36** | 0.034 | 0.001 | 128.751 | cgcg,acag |
| **H75.37** | 0.026 | 0.000 | 127.855 | cgcg,agac |
| **H75.38** | 0.018 | 0.001 | 127.055 | cagu,cgcg |
| **H75.39** | 0.015 | 0.000 | 130.303 | cggu,ccac |
| **H75.40** | 0.014 | 0.001 | 134.727 | ggcg,uaaa |
| **H75.41** | 0.013 | 0.001 | 133.192 | ugug,acac |
| **H75.42** | 0.012 | 0.001 | 131.646 | cggu,cgaa |
| **H75.43** | 0.007 | 0.000 | 134.992 | cggu,caac |
| **H75.44** | 0.006 | 0.000 | 131.166 | cgcg,acaa |
| **H75.45** | 0.004 | 0.000 | 136.609 | ccaa,aacc |
| **H75.46** | 0.001 | 0.000 | 132.792 | cgcg,acac |
| **H75.47** | -0.002 | 0.000 | 133.290 | cagu,cgcc |
| **H75.48** | -0.004 | 0.000 | 135.088 | guaa,agcg |
| **H75.49** | -0.006 | 0.002 | 137.776 | caca,ucac |
| **H75.50** | -0.007 | 0.001 | 137.197 | cucg,uccc |

**Table S2. Primers used to build randomized H75 plasmids.**

| | |
|---|---|
| **Insert FP** | TGAACCTTTACTATAGCTTGBDBDTGAACATTGAGCCTTGATGTGT |
| **Insert RP** | CCAGTCAAACTACCCACCAGBDBDTGTCCGCAACCCGGATTA |
| **Backbone FP** | TGGTGGGTAGTTTGACTGGGG |
| **Backbone RP** | TGGTGGGTAGTTTGACTGGGG |

| Construct Name | Relative Activity | Standard Deviation | 2046-2050 | 2618-2622 | SWM Score |
|---|---|---|---|---|---|
| **H73-1** | 1.333 | 0.059 | cgccg | cggcg | -50.677 |
| **H73-2** | 1.261 | 0.045 | cgcag | cugcg | -54.276 |
| **H73-3** | 1.121 | 0.041 | agccu | aggcc | -53.670 |
| **H73-4** | 1.037 | 0.027 | cgcca | cggcg | -55.429 |
| **H73-5** | 0.998 | 0.028 | cgccg | cggcc | -47.495 |
| **H73-6** | 0.941 | 0.029 | caaua | cguug | -40.547 |
| **H73-7** | 0.715 | 0.028 | cgcag | cugug | -54.779 |
| **H73-8** | 0.686 | 0.033 | cgccg | cagcg | -55.794 |
| **H73-9** | 0.659 | 0.028 | agcag | cggca | -38.793 |
| **H73-10** | 0.643 | 0.020 | agccg | aggcc | -40.986 |
| **H73-11** | 0.435 | 0.018 | aguca | agacc | -48.076 |
| **H73-12** | 0.348 | 0.027 | agcug | cagua | -48.787 |
| **H73-13** | 0.314 | 0.011 | cuccg | aggag | -48.627 |
| **H73-14** | 0.082 | 0.005 | agcca | cugca | -48.101 |

| Construct Name | Relative Activity | Standard Deviation | 2523-2527 | 2536-2540 | SWM Score |
|---|---|---|---|---|---|
| **H91-1** | 1.178 | 0.036 | ccuag | cgggg | -32.745 |
| **H91-2** | 1.130 | 0.062 | caccg | cggug | -33.896 |
| **H91-3** | 0.880 | 0.058 | acccg | ccggu | -33.291 |
| **H91-4** | 0.709 | 0.016 | accag | ccggu | -30.450 |
| **H91-5** | 0.573 | 0.022 | cacag | ccggg | -30.475 |
| **H91-6** | 0.537 | 0.042 | ccccg | cgggg | -28.604 |
| **H91-7** | 0.162 | 0.012 | ccacg | ccggg | -31.576 |
| **H91-8** | 0.024 | 0.002 | cucug | aagaa | -32.275 |
| **H91-9** | 0.017 | 0.001 | aauag | ccagu | -32.944 |
| **H91-10** | 0.010 | 0.001 | ccaaa | uaaga | -30.967 |
| **H91-11** | 0.009 | 0.002 | acccu | ugaug | -27.221 |
| **H91-12** | 0.008 | 0.002 | aacau | cgggg | -27.104 |
| **H91-13** | 0.005 | 0.001 | accaa | cggga | -28.561 |
| **H91-14** | 0.005 | 0.000 | ccacg | cagug | -32.170 |

| Construct Name | Relative Activity | Standard Deviation | 2547-2551 | 2557-2561 | SWM Score |
|---|---|---|---|---|---|
| **H92-1** | 0.344 | 0.002 | ggccg | cgggc | -10.061 |
| **H92-2** | 0.261 | 0.003 | gguca | uggcc | -8.448 |
| **H92-3** | 0.204 | 0.007 | ggacg | cggcc | -6.888 |
| **H92-4** | 0.115 | 0.005 | gccaa | uuggc | -6.631 |
| **H92-5** | 0.072 | 0.003 | gccaa | agggc | -7.065 |
| **H92-6** | 0.058 | 0.001 | cgcca | aggcg | -11.034 |
| **H92-7** | 0.031 | 0.002 | ggcca | cggcc | -9.743 |
| **H92-8** | 0.022 | 0.001 | gcucg | agagc | -10.205 |
| **H92-9** | 0.008 | 0.000 | ggccg | agggc | -8.210 |
| **H92-10** | 0.006 | 0.000 | gccca | cggcc | -8.778 |
| **H92-11** | -0.003 | 0.003 | gccca | agggc | -7.389 |
| **H92-12** | -0.003 | 0.001 | ggcca | agauc | -7.392 |
| **H92-13** | -0.004 | 0.001 | gcaaa | aaaca | -7.494 |

| | | | | | |
|---|---|---|---|---|---|
| **H92-14** | -0.008 | 0.000 | gcaca | cgagg | -7.942 |

**Table S3. SWM scores and sequences of H73, H91 and H92 design constructs.** Sequences and scores of the 14 constructs selected from each design simulation.

**Table S4. Overall SWM score ranges for each library simulation.**

| H73 Design Results | |
|---|---|
| Min Score | -55.794 |
| Max Score | -8.414 |
| Range | 64.208 |

| H91 Design Results | |
|---|---|
| Min Score | -33.896 |
| Max Score | 12.121 |
| Range | 46.017 |

| H92 Design Results | |
|---|---|
| Min Score | -11.034 |
| Max Score | 15.442 |
| Range | -26.476 |

**Table S5. Sequences and iSAT activities of all single and multi-mutants.** "Activity" is signal of construct in iSAT relative to the wildtype ribosome's signal.

| Name | Mutant Combination | H73 Sequence | H75 Sequence | H91 Sequence | H92 Sequence | Activity | Std. Dev. |
|---|---|---|---|---|---|---|---|
| **C1** | 73.1,75.3,91.2,92.1 | cgccg, cggcg | guuu, aacc | caccg, cggug | ggccg, cgggc | -0.003 | 0.001 |
| **C2** | 73.1,75.3,91.2,H92-WT | cgccg, cggcg | guuu, aacc | caccg, cggug | atggc, gccat | 0.485 | 0.007 |
| **C3** | 73.1,75.3,91.1,92.1 | cgccg, cggcg | guuu, aacc | ccuag, cgggg | ggccg, cgggc | -0.005 | 0.001 |
| **C4** | 73.1,75.3,91.1,H92-WT | cgccg, cggcg | guuu, aacc | ccuag, cgggg | atggc, gccat | 0.228 | 0.066 |
| **C5** | 73.1,75.3,H91-WT,92.1 | cgccg, cggcg | guuu, aacc | ggggc, gtccc | ggccg, cgggc | 0.010 | 0.000 |
| **C6** | 73.1,75.3,H91-WT,H92-WT | cgccg, cggcg | guuu, aacc | ggggc, gtccc | atggc, gccat | 0.397 | 0.021 |
| **C7** | 73.1,75.1,91.2,92.1 | cgccg, cggcg | gguu, cacc | caccg, cggug | ggccg, cgggc | -0.001 | 0.000 |
| **C8** | 73.1,75.1,91.2,H92-WT | cgccg, cggcg | gguu, cacc | caccg, cggug | atggc, gccat | 0.253 | 0.030 |
| **C9** | 73.1,75.1,91.1,92.1 | cgccg, cggcg | gguu, cacc | ccuag, cgggg | ggccg, cgggc | -0.001 | 0.001 |
| **C10** | 73.1,75.1,91.1,H92-WT | cgccg, cggcg | gguu, cacc | ccuag, cgggg | atggc, gccat | 0.651 | 0.033 |
| **C11** | 73.1,75.1,H91-WT,92.1 | cgccg, cggcg | gguu, cacc | ggggc, gtccc | ggccg, cgggc | 0.038 | 0.006 |
| **C12** | 73.1,75.1,H91-WT,H92-WT | cgccg, cggcg | gguu, cacc | ggggc, gtccc | atggc, gccat | 0.854 | 0.070 |
| **C13** | 73.1,75.2,91.2,92.1 | cgccg, cggcg | cggg, cccg | caccg, cggug | ggccg, cgggc | -0.004 | 0.001 |
| **C14** | 73.1,75.2,91.2,H92-WT | cgccg, cggcg | cggg, cccg | caccg, cggug | atggc, gccat | 0.055 | 0.016 |
| **C15** | 73.1,75.2,91.1,92.1 | cgccg, cggcg | cggg, cccg | ccuag, cgggg | ggccg, cgggc | 0.004 | 0.005 |
| **C16** | 73.1,75.2,91.1,H92-WT | cgccg, cggcg | cggg, cccg | ccuag, cgggg | atggc, gccat | 0.067 | 0.014 |
| **C17** | 73.1,75.2,H91-WT,92.1 | cgccg, cggcg | cggg, cccg | ggggc, gtccc | ggccg, cgggc | 0.002 | 0.002 |
| **C18** | 73.1,75.2,H91-WT,H92-WT | cgccg, cggcg | cggg, cccg | ggggc, gtccc | atggc, gccat | 0.741 | 0.057 |
| **C19** | 73.1,H75-WT,91.2,92.1 | cgccg, cggcg | acac, gugu | caccg, cggug | ggccg, cgggc | -0.003 | 0.000 |
| **C20** | 73.1,H75-WT,91.2,H92-WT | cgccg, cggcg | acac, gugu | caccg, cggug | atggc, gccat | 0.350 | 0.033 |
| **C21** | 73.1,H75-WT,91.1,92.1 | cgccg, cggcg | acac, gugu | ccuag, cgggg | ggccg, cgggc | -0.006 | 0.001 |
| **C22** | 73.1,H75-WT,91.1,H92-WT | cgccg, cggcg | acac, gugu | ccuag, cgggg | atggc, gccat | 0.328 | 0.046 |
| **C23** | 73.1,H75-WT,H91-WT,92.1 | cgccg, cggcg | acac, gugu | ggggc, gtccc | ggccg, cgggc | 0.000 | 0.003 |
| **C24** | 73.2,75.3,91.2,92.1 | cgcag, cugcg | guuu, aacc | caccg, cggug | ggccg, cgggc | -0.005 | 0.001 |
| **C25** | 73.2,75.3,91.2,H92-WT | cgcag, cugcg | guuu, aacc | caccg, cggug | atggc, gccat | 0.352 | 0.003 |
| **C26** | 73.2,75.3,91.1,92.1 | cgcag, cugcg | guuu, aacc | ccuag, cgggg | ggccg, cgggc | -0.003 | 0.000 |
| **C27** | 73.2,75.3,91.1,H92-WT | cgcag, cugcg | guuu, aacc | ccuag, cgggg | atggc, gccat | 0.262 | 0.003 |
| **C28** | 73.2,75.3,H91-WT,92.1 | cgcag, cugcg | guuu, aacc | ggggc, gtccc | ggccg, cgggc | -0.001 | 0.000 |
| **C29** | 73.2,75.3,H91-WT,H92-WT | cgcag, cugcg | guuu, aacc | ggggc, gtccc | atggc, gccat | 0.845 | 0.044 |
| **C30** | 73.2,75.1,91.2,92.1 | cgcag, cugcg | gguu, cacc | caccg, cggug | ggccg, cgggc | -0.001 | 0.000 |
| **C31** | 73.2,75.1,91.2,H92-WT | cgcag, cugcg | gguu, cacc | caccg, cggug | atggc, gccat | 0.484 | 0.027 |
| **C32** | 73.2,75.1,91.1,92.1 | cgcag, cugcg | gguu, cacc | ccuag, cgggg | ggccg, cgggc | -0.001 | 0.000 |
| **C33** | 73.2,75.1,91.1,H92-WT | cgcag, cugcg | gguu, cacc | ccuag, cgggg | atggc, gccat | 0.450 | 0.019 |
| **C34** | 73.2,75.1,H91-WT,92.1 | cgcag, cugcg | gguu, cacc | ggggc, gtccc | ggccg, cgggc | 0.028 | 0.001 |
| **C35** | 73.2,75.1,H91-WT,H92-WT | cgcag, cugcg | gguu, cacc | ggggc, gtccc | atggc, gccat | 1.243 | 0.023 |
| **C36** | 73.2,75.2,91.2,92.1 | cgcag, cugcg | cggg, cccg | caccg, cggug | ggccg, cgggc | 0.003 | 0.001 |
| **C37** | 73.2,75.2,91.2,H92-WT | cgcag, cugcg | cggg, cccg | caccg, cggug | atggc, gccat | 0.431 | 0.039 |
| **C38** | 73.2,75.2,91.1,92.1 | cgcag, cugcg | cggg, cccg | ccuag, cgggg | ggccg, cgggc | 0.001 | 0.001 |
| **C39** | 73.2,75.2,91.1,H92-WT | cgcag, cugcg | cggg, cccg | ccuag, cgggg | atggc, gccat | 0.035 | 0.006 |
| **C40** | 73.2,75.2,H91-WT,92.1 | cgcag, cugcg | cggg, cccg | ggggc, gtccc | ggccg, cgggc | 0.006 | 0.002 |
| **C41** | 73.2,75.2,H91-WT,H92-WT | cgcag, cugcg | cggg, cccg | ggggc, gtccc | atggc, gccat | 0.340 | 0.016 |
| **C42** | 73.2,H75-WT,91.2,92.1 | cgcag, cugcg | acac, gugu | caccg, cggug | ggccg, cgggc | 0.001 | 0.001 |
| **C43** | 73.2,H75-WT,91.2,H92-WT | cgcag, cugcg | acac, gugu | caccg, cggug | atggc, gccat | 0.001 | 0.001 |
| **C44** | 73.2,H75-WT,91.1,92.1 | cgcag, cugcg | acac, gugu | ccuag, cgggg | ggccg, cgggc | 0.419 | 0.029 |
| **C45** | 73.2,H75-WT,91.1,H92-WT | cgcag, cugcg | acac, gugu | ccuag, cgggg | atggc, gccat | 0.454 | 0.051 |
| **C46** | 73.2,H75-WT,H91-WT,92.1 | cgcag, cugcg | acac, gugu | ggggc, gtccc | ggccg, cgggc | 0.010 | 0.002 |
| **C47** | H73-WT,75.3,91.2,92.1 | gcggc, gccgt | guuu, aacc | caccg, cggug | ggccg, cgggc | 0.000 | 0.001 |
| **C48** | H73-WT,75.3,91.2,H92-WT | gcggc, gccgt | guuu, aacc | caccg, cggug | atggc, gccat | 1.148 | 0.025 |
| **C49** | H73-WT,75.3,91.1,92.1 | gcggc, gccgt | guuu, aacc | ccuag, cgggg | ggccg, cgggc | -0.001 | 0.000 |
| **C50** | H73-WT,75.3,91.1,H92-WT | gcggc, gccgt | guuu, aacc | ccuag, cgggg | atggc, gccat | 0.420 | 0.026 |
| **C51** | H73-WT,75.3,H91-WT,92.1 | gcggc, gccgt | guuu, aacc | ggggc, gtccc | ggccg, cgggc | 0.051 | 0.003 |
| **C52** | H73-WT,75.1,91.2,92.1 | gcggc, gccgt | gguu, cacc | caccg, cggug | ggccg, cgggc | 0.005 | 0.001 |
| **C53** | H73-WT,75.1,91.2,H92-WT | gcggc, gccgt | gguu, cacc | caccg, cggug | atggc, gccat | 1.232 | 0.006 |
| **C54** | H73-WT,75.1,91.1,92.1 | gcggc, gccgt | gguu, cacc | ccuag, cgggg | ggccg, cgggc | 0.007 | 0.002 |

| | | | | | | | |
|---|---|---|---|---|---|---|---|
| **C55** | H73-WT,75.1,91.1,H92-WT | gcggc, gccgt | gguu, cacc | ccuag, cgggg | atggc, gccat | 0.410 | 0.022 |
| **C56** | H73-WT,75.1,H91-WT,92.1 | gcggc, gccgt | gguu, cacc | ggggc, gtccc | ggccg, cgggc | 0.179 | 0.002 |
| **C57** | H73-WT,75.2,91.2,92.1 | gcggc, gccgt | cggg, cccg | caccg, cggug | ggccg, cgggc | 0.005 | 0.000 |
| **C58** | H73-WT,75.2,91.2,H92-WT | gcggc, gccgt | cggg, cccg | caccg, cggug | atggc, gccat | 0.004 | 0.005 |
| **C59** | H73-WT,75.2,91.1,92.1 | gcggc, gccgt | cggg, cccg | ccuag, cgggg | ggccg, cgggc | 0.036 | 0.003 |
| **C60** | H73-WT,75.2,91.1,H92-WT | gcggc, gccgt | cggg, cccg | ccuag, cgggg | atggc, gccat | 0.542 | 0.023 |
| **C61** | H73-WT,75.2,H91-WT,92.1 | gcggc, gccgt | cggg, cccg | ggggc, gtccc | ggccg, cgggc | 0.089 | 0.010 |
| **C62** | H73-WT,H75-WT,91.2,92.1 | gcggc, gccgt | acac, gugu | caccg, cggug | ggccg, cgggc | 0.012 | 0.000 |
| **C63** | H73-WT,H75-WT,91.1,92.1 | gcggc, gccgt | acac, gugu | ccuag, cgggg | ggccg, cgggc | 0.101 | 0.008 |
| **H92.1** | H73-WT,H75-WT,H91-WT,H92-7 | gcggc, gccgt | acac, gugu | ggggc, gtccc | ggccg, cgggc | 0.344 | 0.002 |
| **H91.1** | H73-WT,H75-WT,H91-11,H92-WT | gcggc, gccgt | acac, gugu | ccuag, cgggg | atggc, gccat | 1.178 | 0.036 |
| **H91.2** | H73-WT,H75-WT,H91-9,H92-WT | gcggc, gccgt | acac, gugu | caccg, cggug | atggc, gccat | 1.130 | 0.062 |
| **H73.1** | H73-10,H75-WT,H91-WT,H92-WT | cgccg, cggcg | acac, gugu | ggggc, gtccc | atggc, gccat | 1.333 | 0.059 |
| **H75.2** | H73-WT,H75-39,H91-WT,H92-WT | gcggc, gccgt | cggg, cccg | ggggc, gtccc | atggc, gccat | 1.387 | 0.013 |
| **H73.2** | H73-8,H75-WT,H91-WT,H92-WT | cgcag, cugcg | acac, gugu | ggggc, gtccc | atggc, gccat | 1.261 | 0.045 |
| **H75.3** | H73-WT,H75-43,H91-WT,H92-WT | gcggc, gccgt | guuu, aacc | ggggc, gtccc | atggc, gccat | 1.394 | 0.028 |
| **H75.1** | H73-WT,H75-41,H91-WT,H92-WT | gcggc, gccgt | gguu, cacc | ggggc, gtccc | atggc, gccat | 1.400 | 0.022 |
| **Wildtype** | H73-WT,H75-WT,H91-WT,H92-WT | gcggc, gccgt | acac, gugu | ggggc, gtccc | atggc, gccat | 0.979 | 0.029 |

#### Supplementary Figures

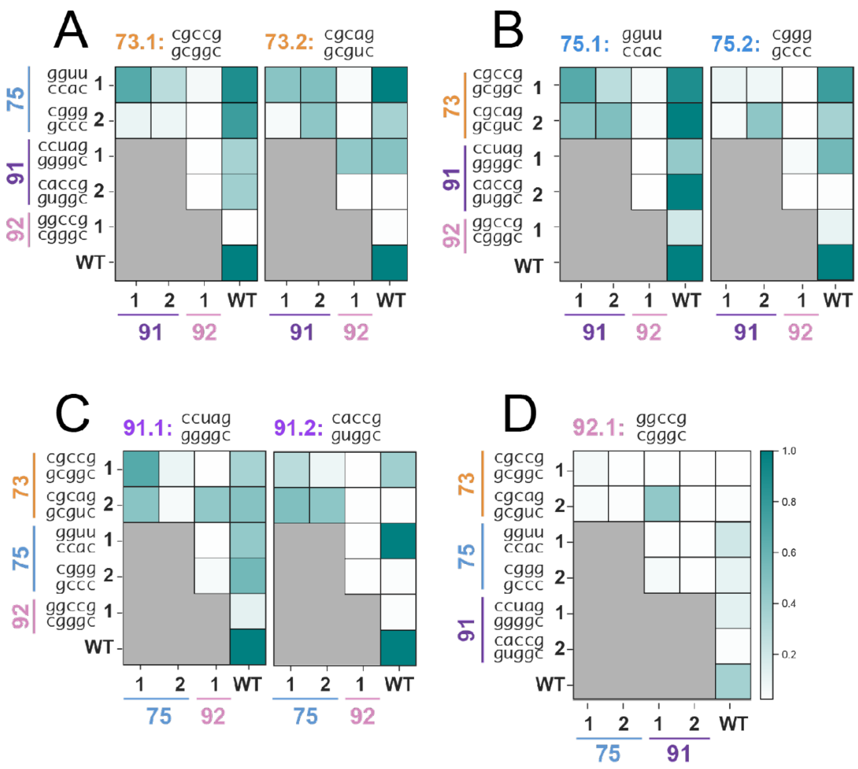


**Figure S1. Complete heatmaps of combination construct activities in iSAT.**
